# The geometry of gametic dispersal in a flying mammal, *Rhinolophus hipposideros*

**DOI:** 10.1101/2024.10.24.620000

**Authors:** Thomas Brazier, Diane Zarzoso-Lacoste, Lisa Lehnen, Pierre-Loup Jan, Sebastien J. Puechmaille, Eric J. Petit

## Abstract

Dispersal influences population and evolutionary dynamics, in a manner that depends on by which dispersal strategies gene flow occurs. In some species, mating partners move exclusively for mating, dispersing genes but not individuals. This is the case in many bat species, of which the lesser horseshoe bat (*Rhinolophus hipposideros*) shows a genetic structure at a fine spatial scale suggesting restricted dispersal. We investigated how natal and mating dispersal shape gene flow in this species in two metapopulations using paternity and population assignments. Half of the inferred paternities were intra-colonial and gave an estimate of the mean mating dispersal distance of around 11 km, explaining the observed genetic structure. Complete gametic dispersal distances were further estimated by combining natal with mating dispersal distances. The resulting gametic dispersal kernels showed a mean distance of around 20 km and a fat-tailed distribution typical of an excess of long-distance dispersal movements. It is the first time that natal and mating dispersal distances have been separately estimated and then combined in animals, documenting quantitatively how mating dispersal decorrelates gene and individual flows. It is important to consider this mechanism to explain dispersal evolution.

## Introduction

Individuals or propagules may move between their birthplace or the place where they are produced and any other place, habitat, social group, patch, etc. These movements, called dispersal, change the composition of the community they leave and the community they settle in, modify the demography and trait distribution of both the population they leave and the population they settle in, and exchange genes between populations. Dispersal is involved in range expansion and colonizations, the rescue of small populations from extinction, invasiveness of exotic species but also the spread of pathogens or diseases, and it affects source-sink dynamics (Brown and Kodric-Brown, 1977; Gadgil, 1971; Hastings et al., 2004; Perkins et al., 2013; Phillips et al., 2008). In addition, dispersing individuals contribute to gene flow, affecting local adaptation or the spread of beneficial mutations, for example (Peischl and Gilbert, 2020; Tigano and Friesen, 2016). Dispersal is thus of crucial importance for both ecology and evolution, since population and evolutionary dynamics are intertwined through, among others, this particular trait.

Dispersal strategies are largely diversified among species (Bowler and Benton, 2005), mostly depending on proportions of dispersing individuals, mobility, and life-stage involved (Broquet and Petit, 2009). Natal dispersal, the definitive movement of propagules outside their natal location, generally occurs before the first reproduction event. Breeding dispersal is a later temporary movement that occurs between successive reproductive bouts and locations. Mating dispersal, which is relative to the pairing of opposite-sex individuals from different resident sites aimed for reproduction, also occurs after natal dispersal (Hazlitt et al., 2006; Sugg et al., 1996). Breeding and mating dispersal are therefore two alternative dispersal strategies that occur after natal dispersal. To our knowledge, there are no documented cases of species exhibiting all three types of dispersal, meaning that a full understanding of dispersal and its consequences should consider at least two of these strategies when either breeding or mating dispersal occurs. This has been demonstrated in birds, where breeding dispersal is widespread (Dale et al., 2005; Fandos et al., 2023; Paradis et al., 1998). Dispersal can be gametic and/or zygotic, depending on which propagules are involved in movements (gametes vs. individuals). This is most easily illustrated in plants, in which seed dispersal is zygotic while pollen dispersal is gametic. Natal dispersal can be either gametic or zygotic (see the example with plants). Breeding dispersal is only zygotic, as it involves the permanent movement of adult individuals. In contrast, mating dispersal does not involve a permanent movement of parents and leads to gametic dispersal only. Therefore, natal, breeding, and mating dispersal contribute to gene flow and the dispersal of individuals to different degrees (Waser and Elliott, 1991).

Importantly, male gametic dispersal may result from the addition of different dispersal behaviours: in mobile species, the addition of natal dispersal movements to mating dispersal movements results in male gametes moving further than the male home site. Natal dispersal is assessable by multiple methods (e.g. Capture-Mark-Recapture, direct observation or tagging). Mating dispersal, in contrast, involves home site fidelity and only short (sometimes a few hours), temporary movement to another site and is thus difficult to assess. Consequently, the contribution of mating dispersal to gene flow may be underestimated, because movements for mating often remain cryptic. Yet, mating dispersal has been documented in invertebrates, fishes, lizards, birds and mammals (Archie et al., 2008; Chapple and Keogh, 2005; Johannesen and Lubin, 1999; Kesler et al., 2010; Portnoy et al., 2015).

The lesser horseshoe bat, *Rhinolophus hipposideros* (André, 1797), is an ideal species to study natal and mating dispersal separately. Like most temperate bat species, it is sedentary and females are philopatric. Dispersal is thus supported by males (Bontadina et al., 2002; Gaisler and Chytil, 2002). As in many other bat species, little is known about the mating system and male dispersal (McCracken and Wilkinson, 2000). Because male-biased dispersal is pervasive among mammals (Dobson, 1982; Greenwood, 1980; Lawson Handley and Perrin, 2007), including bats (Kerth et al., 2002; Petit et al., 2001), it is assumed that males actively disperse and participate to gene flow through natal dispersal and mating with females that are not from their natal colony. In such species, natal and mating dispersal movements need to be studied simultaneously to reveal their relative contribution to gametic dispersal and ultimately gene flow (Archie et al., 2008; Chapple and Keogh, 2005; Kerth and Morf, 2004). Interestingly, Lehnen et al. (2021) recently demonstrated that the lesser horseshoe bat’s genetic structure follows an isolation-by-distance pattern at a scale of a few kilometres in two different metapopulations. This pattern is unusual among bats, suggesting very limited gene flow in this species (Aguillon et al., 2017; Rousset, 1997).

We aim to characterize the geometry of male gametic dispersal by following the spatial displacement of sperm through both natal and mating dispersal movements, adopting a dispersal distance kernel perspective (Nathan et al., 2012). We tested the hypotheses that (1) some lesser horseshoe bat males adopt a philopatric behaviour (Entwistle et al., 2000) and (2) males mate preferably with females of their home colony, which is either their natal colony or the colony they settle in after their natal dispersal movement, if any. We used molecular methods to leverage the strong population structure at a fine scale, assigning fathers to their colony of birth for estimating natal dispersal, and assigning offspring to their father to estimate mating dispersal (Broquet and Petit, 2009; Dussex et al., 2016; Pemberton, 2008). Natal and mating dispersal distances were then combined in a continuous-space model to estimate the complete male gametic dispersal distance kernel.

## Materials and methods

### Study metapopulations

We analysed data previously gathered on two lesser horseshoe bat metapopulations with different population dynamics (Jan et al., 2019; Lehnen et al., 2021; Zarzoso-Lacoste et al., 2018). The French metapopulation (Picardy) was stable while the German one (Thuringia) was expanding. Seventeen colonies were sampled over four years (2013-2016) in Picardy (France), and 22 colonies were sampled over three years (2015-2017) in Thuringia (Germany). We excluded from the data set in Zarzoso-Lacoste et al. (2018) one remote colony isolated in a forest that is not located in our study area and a second one that we were not able to sample repeatedly across years. Distances between colonies ranged from 0.7 to 56 km in Picardy and from 1.8 to 99 km in Thuringia (Fig. S1).

### Genotype dataset

The dataset comprised all unique genotypes obtained from bats faeces sampled in the colonies during the respective study periods. Each unique genotype of six to eight microsatellite markers was considered as one individual (see Zarzoso-Lacoste et al. 2018, 2020 for the complete procedure). There were two sampling sessions per year, during summer, before and after parturition. All individuals sampled for the first time after parturition were considered as putative offspring. Females and males were separated by the sexual marker DDX3X/Y-Mam with the method described by Zarzoso-Lacoste et al. (2018, 2020). Summary statistics for the microsatellite dataset were estimated with the R packages adegenet and hierfstat (Goudet and Jombart, 2022; Jombart, 2008; Jombart and Ahmed, 2011).

### Parentage assignment

Maternities and paternities were inferred with COLONY 2 (Jones and Wang, 2010) version 2.0.6.4 for Linux. COLONY was run with a full-pedigree likelihood method. Following Foley et al. (2020), parental links were inferred from 30 independent COLONY runs of the same dataset with different random seeds to achieve convergence on the best putative father. The eight microsatellite markers were used in the parentage assignment procedure. As COLONY is able to accommodate for genotyping errors, allelic dropout rates and other typing errors at each locus were set with values estimated by Zarzoso-Lacoste et al. (2018). Females were considered as polygamous with only one offspring per year (the father could change every year) and males as polygamous with no offspring limitation. Detection probabilities for mothers and fathers were set to 0.9 for mothers and 0.56 for fathers (details in Supplementary Information). Cohorts of the different years (four in Picardy, three in Thuringia) were analysed together.

In this species, both sexes are sexually mature at the end of their first year (Gaisler, 1966). Thus, putative offspring were all individuals sampled for the first time after parturition in any given year and these potential offspring were assumed to become candidate parents the year after their birth. Candidate parents were assumed to breed every year. As a consequence, for a given offspring, candidate mothers (respectively fathers) were all females (respectively males) born before the year of sampling of the offspring.

Then, we built lists of excluded paternities and maternities (Jones and Wang, 2010). Excluded paternities were all males born the same or later years for each given offspring. Excluded maternities were also all females born the same or later years for each given offspring, with an additional exclusion of all females from a different colony than the offspring colony as offsprings stay in their mother’s colony for at least several weeks after birth. Females give birth to at most one offspring per year (Gaisler, 1966), so all offspring born the same year as the focal one were excluded from maternal sibship.

As in Foley et al. (2020), the comparison of simulated parent/offspring genotypes and the parentage inferred by COLONY was used both to find a statistic discriminant that allowed to exclude wrong paternities and to perform a power analysis of the categorical assignment method (i.e. type I and type II error rates). We simulated genotypes sampled in a theoretical population with known parent-offspring relationships and used it as an input in COLONY to assess type I (the wrong parent was inferred) and type II (no parent was found among the sampled ones) errors. We simulated the genotype datasets with the COLONY simulation module for Windows version 2.0.6.4 (Wang, 2013) to mimic demography, genetic diversity and genotyping errors observed in each sampled metapopulation (details in supplementary materials and methods). Each simulated dataset was replicated 50 times with different random seeds then analysed with COLONY to assess for a convergence on true fathers over multiple replicated runs.

### Male mating dispersal

The empirical distribution of mating dispersal distances was directly assessed by pairwise distances between the home colony of offspring and fathers inferred with categorical assignment (i.e. COLONY). We compared these estimates to expected values obtained by a random association between true juveniles (candidate juveniles for which a true parent was found) and candidate fathers. A 95% confidence envelope for this null expectation was obtained by using a bootstrap procedure. The expected and observed distributions of offspring-father distances were compared with a Kolmogorov-Smirnov test. Besides, the exponential parameter *β* of the mating dispersal distance kernel was estimated by a Bayesian approach provided in the MasterBayes package (Hadfield et al., 2006) for cross-validation.

### Male natal dispersal

The natal colony of male individuals was assessed by genetic assignment of fathers to colonies in the dataset with STRUCTURE (Pritchard et al., 2000). Male adults were assigned to colonies based on the genetic structure inferred from all female individuals with the USEPOPINFO model, as described in Dussex et al. (2016). Natal dispersal was then estimated by calculating the distance between the source colony inferred by STRUCTURE and the colony where the male was sampled. Only migrants of the actual generation were searched (GENSBACK = 0). The colony with the highest assignment probability was retained as the source colony for migrants. Ten replicated STRUCTURE analyses were run with a total of 2,000,000 iterations including a burnin of 1,000,000 (thinning = 100). Within-chain convergence was assessed with trace plots and diagnostics implemented in the CODA R package (Plummer et al., 2006). Replicates were compared and the replicate minimizing the Geweke statistics for both Alpha and Log-Likelihood sampling chains was retained for categorical assignments.

### Male gametic dispersal

Parameters of the gametic dispersal distance kernel were estimated based on the distribution of empirical distances. Gametic dispersal distances were the result of both natal and mating dispersal; in other words, the distance between the natal colony of the male and the natal colony of its offspring. Competing probabilistic models were fitted to the data with the FitDistRplus package to estimate the parameters of the dispersal kernel with maximum likelihood estimation. Goodness-of-fit were compared based on AICc (Mazerolle, 2020) and BIC criterion (Delignette-Muller and Dutang, 2015). The most classical dispersal distributions (Gaussian, Exponential, Exponential Power, Weibull and Gamma) were compared (Nathan et al., 2012).

Additionally, we investigated hurdle models to deal with the excess of zeroes in dispersal distances (Lachenbruch, 2002). Hurdle models were constructed as a mixture of a binomial distribution for the probability of mating inside/outside of the colony and a continuous distribution excluding null dispersal distances. The parameter of the binomial distribution was estimated with a Generalized Linear Model in R and parameters of the positive continuous distribution were estimated with FitDistRplus. Parameters’ confidence intervals were computed by bootstrapping. The log-likelihood of hurdle models was the sum of the log-likelihoods of the binomial model and the conditional model separately estimated (McDowell, 2003).

### Statistical analysis

All statistical analyses were conducted with R version 4.1.2 (R Core Team, 2022). From the distributions of observed mating and gametic offspring-father distances, we estimated the mean and median dispersal distances with a bootstrap method.

## Results

### Dataset

Of the 3,706 unique genotypes in Picardy (France), 1,602 individuals were labelled as offspring (43%). 72% of individuals considered as adults were females (58% to 92% females depending on colony). 21% of individuals had an incomplete genotype (1 or 2 missing loci). The mean sample size per colony was 218 (SD = 152) individuals for cumulated years 2013-2016, ranging from a minimum sample size of 53 bats to a maximum sample size of 746 bats. In Thuringia (Germany), there were 3,913 unique genotypes with 2,128 potential offspring (54%). 62% of individuals considered as adults were females. After removing one colony with only one individual, the mean sample size per colony was 186 for years 2015-2017 (SD = 108, range 24 to 385). The eight microsatellite markers showed a high diversity both in France and Germany, with an averaged observed heterozygosity across loci and populations of 0.61 in Picardy and 0.53 in Thuringia, and a mean allelic richness of 7.3 and 4.3, respectively (Table S1).

### Simulations

The proportion of successful father assignments was low with a high prevalence of type II errors (no father found for a given offspring, Table S2). In simulations, a conservative frequency value of 0.3 in Picardy and 0.25 in Thuringia accurately excluded wrong fathers when at least 20 runs of COLONY were used, even for the least powerful analysis with 6/8 complete loci (Fig. 1).

**Figure 1:**
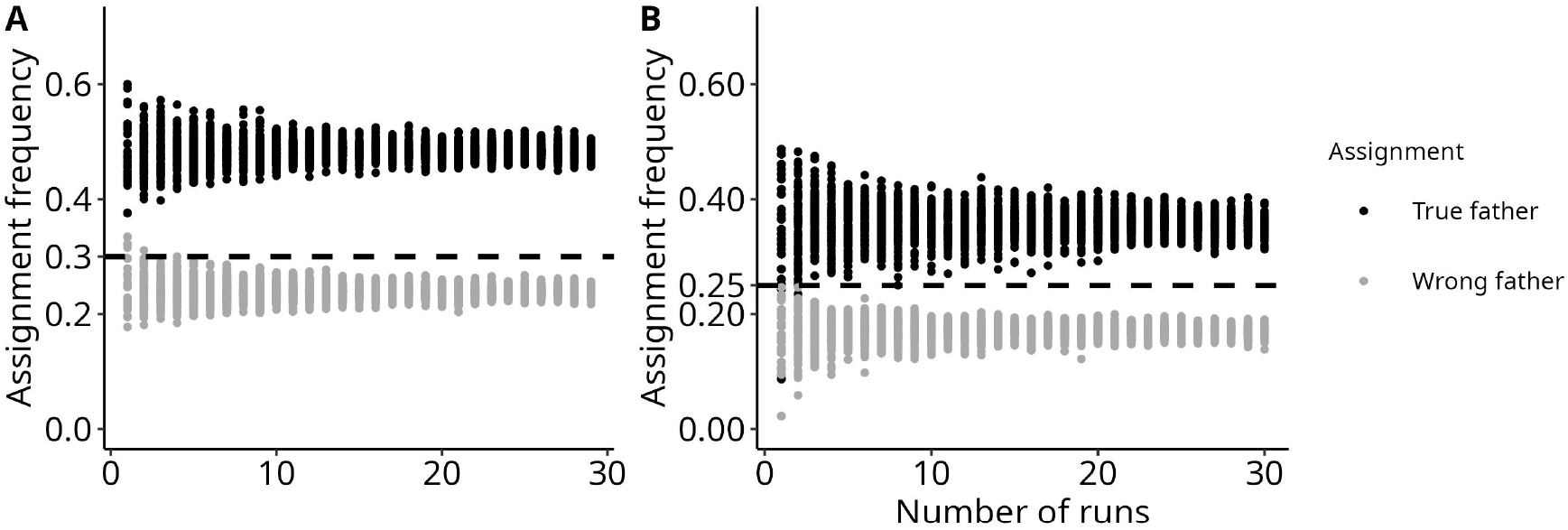
Assignment frequency of the inferred father according to the number of runs in simulated datasets with missing data (6/8 complete genotypes). Black dots indicate cases where a true parent was found; grey dots indicate cases where the wrong parent was found for 3892 simulated offspring. Assignment probabilities of simulated father/offspring dyads with allelic frequencies of Picardy (A) or Thuringia (B). For each number of runs *R*, 100 assignment frequencies were estimated by randomly sampling *R* runs. The dashed line indicates the exclusion threshold of 0.3 (Picardy) or 0.25 (Thuringia) determined in the simulations.

### Parentage assignment

We pooled inferences from genotypes with no more than 2 missing loci (*≥* 6/8 loci) with the inferences from the complete genotypes (8/8 loci). Ambiguous assignments (different fathers for the same offspring) were resolved in favour of the most powerful dataset (8/8 complete loci). COLONY found 530 unique paternities (33% of putative offspring) in the French dataset. Based on simulation results (Fig. 1), we removed inferred paternities found in less than 30% of 30 runs, resulting in 82 paternities being considered as true paternities. There were 17 fathers responsible for these 82 paternities over four years (Table S3). From the German dataset with the same procedure, 62 paternities were retained with only 17 fathers over three years (Table S4).

### Mating dispersal: where do the fathers come from?

Based on the empirical distribution of mating dispersal distances, the Kolmogorov-Smirnov test between the two distributions of offspring-father distances (expected and observed) showed a significant deviation (D = 0.40 in France and 0.53 in Germany, p *<* 0.001 in both cases, n = 82 and 62) with more zero distances than expected by chance (Fig. 2). The mean mating dispersal distance was 10 km (IC95 = [7.11; 13.1], 10,000 bootstraps, n = 82) in France and 11.6 km (IC95 = [7.3; 16.6], 10,000 bootstraps, n = 62) in Germany. However, the median distance were much lower in these two populations (2.3 km, IC95 = [0, 7.7] and 4.7 km, IC95 = [0, 7.8], respectively). Both empirical distributions were globally congruent (Fig. 2) and the MasterBayes Bayesian approach yielded similar results (Supplementary Information).

**Figure 2:**
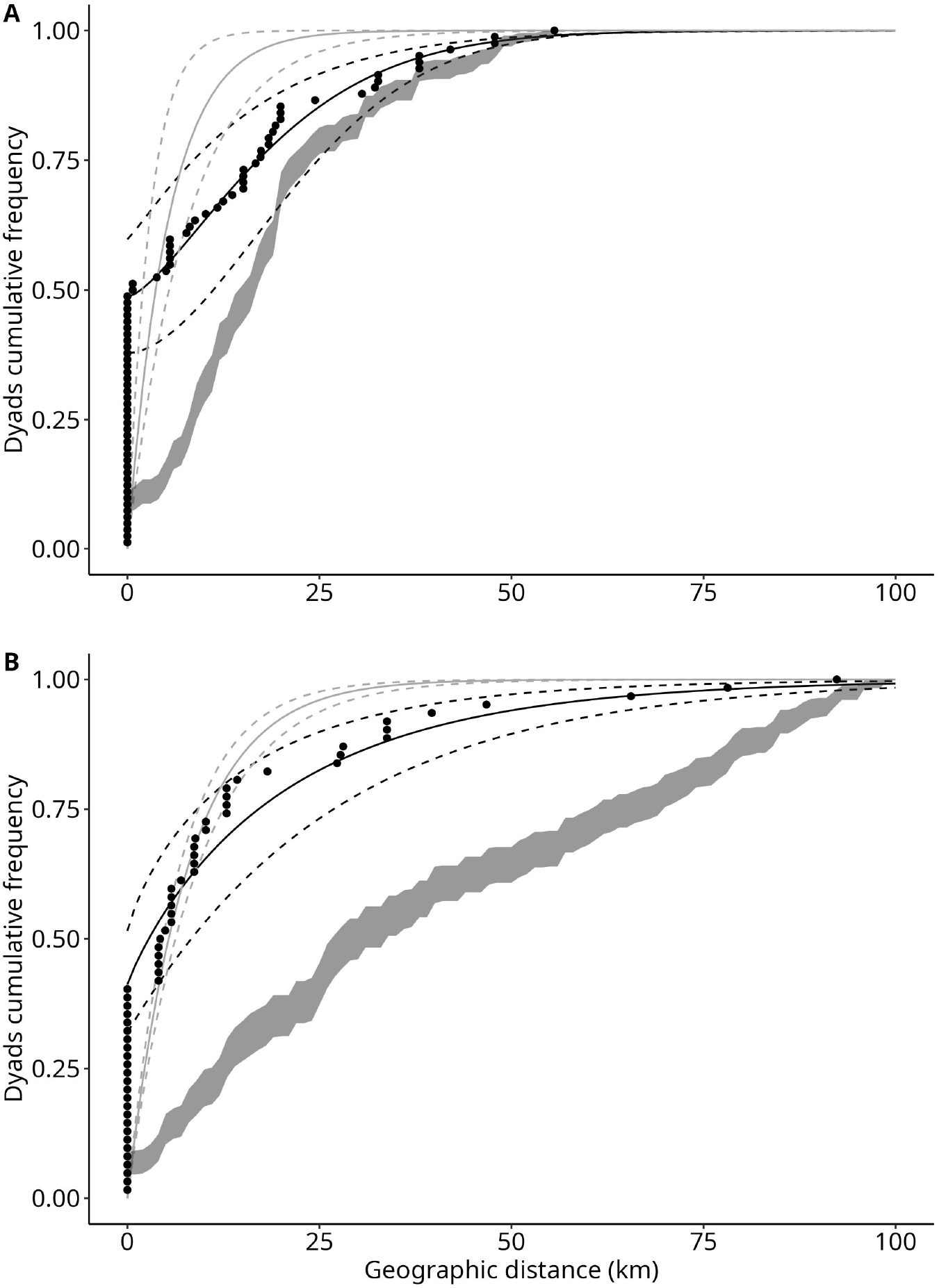
Distribution of offspring-father mating dispersal distances in Picardy (A) and Thuringia (B). Black dots represent observed distances for accepted paternities (n = 82 offspring and 17 fathers in A and n = 62 offspring and 17 fathers in B). The grey ribbon marks the 95% confidence interval for expected distances between candidate fathers and true offspring under a random mating hypothesis, estimated with resampling methods. The black curve shows the theoretical Weibull dispersal kernel (i.e. cumulative density function of distances) with parameters fitted with fitditrplus on individual mating distances (dashed curves correspond to 95% confidence intervals estimated with 1,000 bootstraps). The grey curve shows the negative exponential cumulative density function of distances with the *β* parameter estimated by Masterbayes (dashed curves correspond to 95% confidence intervals estimated with 1,000 bootstraps).

### Natal dispersal: where were the fathers born?

Assignment to natal colonies with STRUCTURE indicated approximately equal proportions of philopatric males and natal dispersers. Of 17 fathers in Picardy, nine were inferred as natal dispersers (52%), yet assignment probabilities were globally low with a mean probability of 0.13. There were 7 dispersing fathers in Thuringia (of a total of 17 fathers, 41%) with a mean assignment probability of 0.52. Assignment probabilities for the inferred fathers in France and Germany were globally among the highest probabilities of the whole dataset (Fig. 3). However, assignment probabilities were globally lower in Picardy than in Thuringia (Fig. 3), with a lower power of discrimination between the best assignment probability and the second best in Picardy (Fig. S2). In contrast, fathers in Germany were easier to assign to a single candidate colony (Fig. S2) and showed the strongest philopatry. Our estimates were robust across ten MCMC sampling chains, systematically inferring the same natal colonies for all fathers with extremely similar probabilities across independent runs of STRUCTURE (Fig. S3). All sampling chains of STRUCTURE globally converged to stationarity and Geweke Z-scores of the selected replicate for Picardy were -0.45 (*α*) and 0.44 (Log-Likelihood). Geweke Z-scores for Thuringia were 0.06 (*α*) and 0.58 (Log-Likelihood).

**Figure 3:**
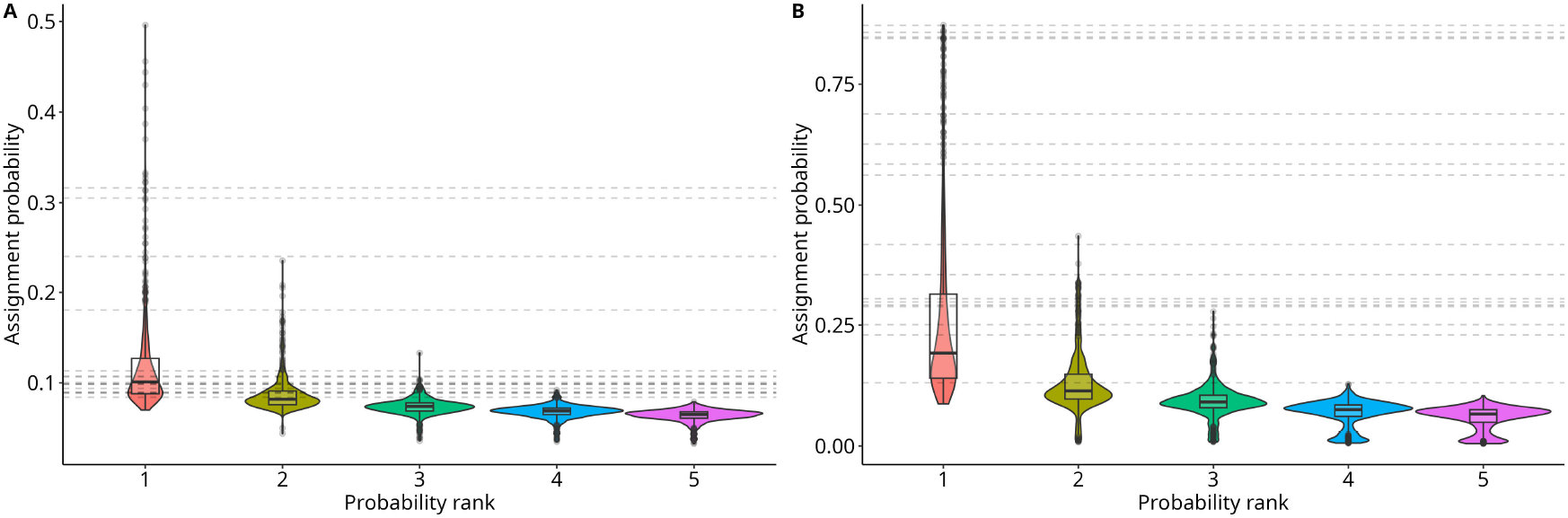
Distributions of assignment probabilities of all male adults to a putative natal colony in France (A) and Germany (B) (n = 578 and 683, respectively). All probabilities estimated with STRUCTURE are represented with different colours, and ordered by decreasing rank of probability values, from the best individual assignment (rank = 1) to the worst (rank = number of colonies). Horizontal dashed lines represent the best probability (rank 1) for fathers previously inferred with COLONY (n = 17 and 17, respectively).

### Gametic dispersal: the whole sperm journey

The mean gametic dispersal distance (i.e. pairwise distance between the natal colony and the colony where the offspring was born resulting from both natal and mating dispersal) was slightly higher than for mating dispersal alone: 17.6 km (IC95 = [14.4; 20.7], 10,000 bootstraps, n = 82) in France and 22.2 km (IC95 = [15.2; 29.6], 10,000 bootstraps, n = 62) in Germany. However, as for mating dispersal, the median gametic dispersal distances were lower than the mean (16.2 km, IC95 = [10.7, 18.9] and 6.9 km, IC95 = [4.1, 8.7], respectively), revealing a strong skewness of the distribution of distances. Upon the individual distances of gametic dispersal, model comparison based on AICc/BIC showed that hurdle models consistently outperformed all single continuous distributions (Table 1). When the conditional probability of mating inside/outside the colony and dispersal distances outside the colony were modelled separately in the hurdle model, the best fitting distribution in Picardy was the binomial + Weibull (Fig. S4). The proportion of gametic dispersal outside the natal colony, estimated by the binomial parameter, was 0.78 (IC95 = [0.68; 0.87], 1,000 bootstraps). The shape and scale parameters of the fitted Weibull were 1.90 (IC95 = [1.54; 2.37], 1,000 bootstraps) and 25.3 (IC95 = [20.9; 29.4], 1,000 bootstraps). In Thuringia, hurdle models outperformed all other ones and Log-Normal was the best fit, yet with very similar AICc/BIC values for different fat-tailed families (Δ_*AICc/BIC*_ *≤* 3; Table 1; Fig. S4). We selected the Weibull model for comparison between Picardy and Thuringia (Fig. 4) and compared it to the negative exponential dispersal kernel inferred by Lehnen et al. (2021) on a large European metapopulation (mean distance = 16.77 km). The binomial parameter in Thuringia was slightly lower than Picardy, at 0.70 (IC95 = [0.56; 0.82], 1,000 bootstraps). The fitted Weibull had a shape of 0.99 (IC95 = [0.87; 1.16], 1,000 bootstraps) and a scale of 31.8 (IC95 = [22.4; 42.5], 1,000 bootstraps).

**Table 1:** Model comparison between competing probabilistic kernels of dispersal distances fitted to the empirical gametic dispersal distribution (i.e. paternity distances inferred with COLONY + natal dispersal inferred with STRUCTURE), based on the number of parameters of the model, AICc and BIC. The best model based on AICc/BIC and all competing models with Δ_*AICc/BIC*_ *≤* 3 are indicated by values in bold.

|  |  | Thuringia |  | Picardy |  |
| --- | --- | --- | --- | --- | --- |
| Model | N parameters | AICc | BIC | AICc | BIC |
| Single continuous distribution |  |  |  |  |  |
| Gaussian | 2 | 599 | 604 | 675 | 679 |
| Negative Exponential | 1 | **510** | **513** | **636** | **638** |
| Exponential power | 3 | 548 | 555 | 747 | 754 |
| Weibull | 2 | - | - | - | - |
| Gamma | 2 | - | - | - | - |
| Hurdle models |  |  |  |  |  |
| Binomial + Gaussian | 3 | 499 | 505 | 597 | 604 |
| Binomial + Negative Exponential | 2 | **465** | **469** | 617 | 622 |
| Binomial + Exponential power | 4 | 480 | 490 | **592** | 603 |
| Binomial + Weibull | 3 | **466** | 473 | **589** | **597** |
| Binomial + Gamma | 3 | **466** | 473 | **592** | **599** |
| Binomial + Log-normal | 3 | **466** | **470** | 601 | 606 |

**Figure 4:**
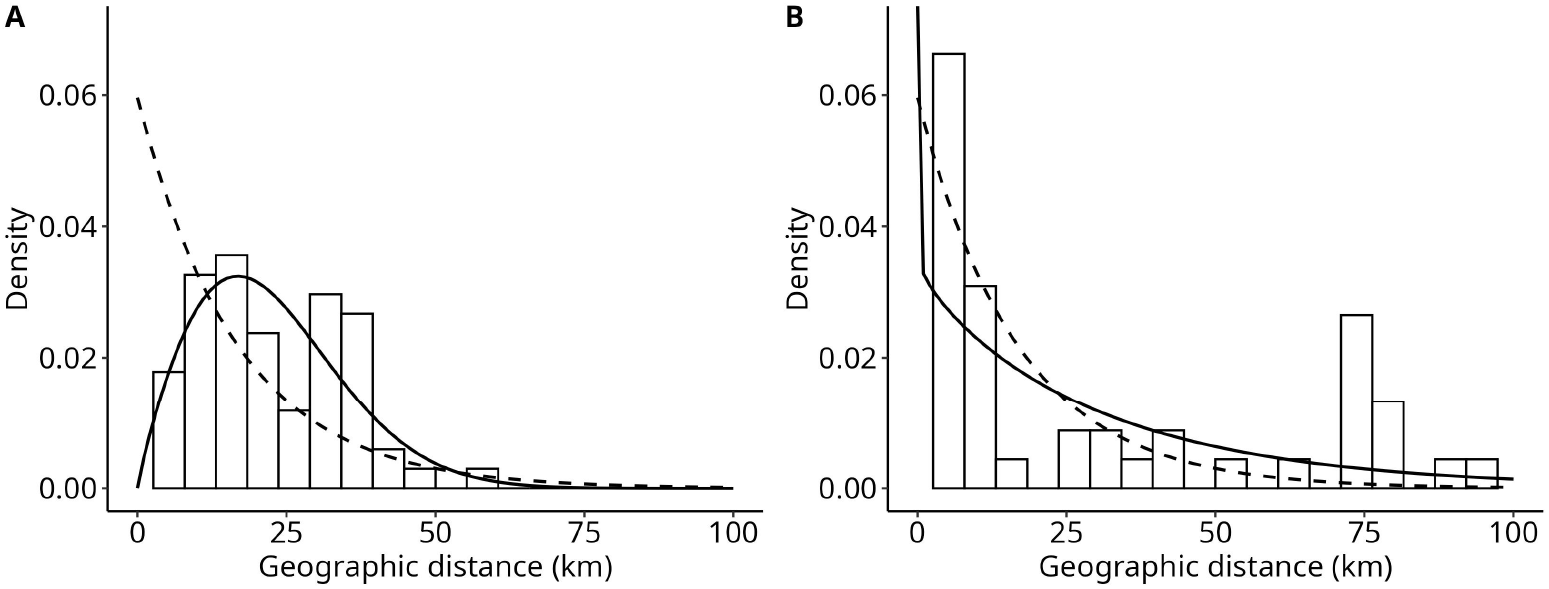
Best model (AICc and BIC criterion) of gametic dispersal distance kernels fitted on strictly positive dispersal distances in Picardy (A) and Thuringia (B). Histograms show densities of empirical gametic dispersal distances. The best model is a Weibull function. In each case, our dispersal kernel (solid line) is compared to the negative exponential dispersal kernel inferred by Lehnen et al. (2021) (dashed line) based on the IBD slope on a large European metapopulation (effective dispersal distance = 16.77 km, shape = 1/16.77).

To assess the effect of potential natal colony mis-assignment on the gametic dispersal kernel, we compared the gametic dispersal kernel inferred with the best assignment probability (positive distances only, Fig. 4) with alternative kernels estimated with all other possible natal colonies, ordered by decreasing assignment probability, i.e. from the most to the least probable natal colony (Fig. S5). By comparing all possible gametic dispersal kernels, from the best to the worst, we showed that the gametic dispersal kernel in France is not qualitatively affected by potential natal colony mis-assignments (Fig. S5A). On the contrary, in Germany, where the power to assign fathers to a natal colony was much higher than in France (Fig. S2), the best kernel was substantially skewed towards short distances compared to kernels with lower assignment probabilities (Fig. S5B).

## Discussion

Both natal and mating dispersal mechanisms contribute to dispersal, and to its impacts on gene, population and community dynamics. These impacts depend on the distribution of dispersal distances, especially when long-distance dispersal is important (Nathan et al., 2012). To the best of our knowledge, our study is the first to consider mating and natal dispersal separately to infer the complete gametic dispersal distance kernel in animals (Nathan, 2007; Rogers et al., 2019).

While mating dispersal has already been explicitly assessed on its own, the contribution of mating dispersal events to gametic dispersal is still scarcely investigated (Archie et al., 2008; Buchalski et al., 2014; Chapple and Keogh, 2005; Randall et al., 2007; Ruch et al., 2009). A previous study reported natal and gametic dispersal distances in a philopatric mammal, the banner-tailed kangaroo rat, with a similar indirect approach as ours, suggesting that mating dispersal contributes most to gene flow in this species, though mating dispersal was not explicitly considered (Winters and Waser, 2003). The combination of pollen and seed dispersal recently received attention in plants (Browne et al., 2018). However, to our knowledge, the de-coupling of mating and natal dispersal to interpret the total gametic dispersal kernel has never been reported in metazoans (Nathan, 2007; Nathan et al., 2012; Rogers et al., 2019).

### Natal and mating dispersal both shape the geometry of the gametic dispersal distance kernel

Dispersal kernels are typically fitted to fat-tailed distributions (i.e. leptokurtic distributions) and most often clearly outperform the Gaussian and Exponential models (Nathan et al., 2012). The gametic dispersal kernel we inferred for the lesser horseshoe bat belongs to this family of models and shows a distribution already observed in many phylogenetically distant species (Browne et al., 2018; Catalano et al., 2020; Collin et al., 2013; Kesler et al., 2010; Paradis et al., 2002). For instance, fat-tailed dispersal kernels are common amongst plant and bird studies (Austerlitz et al., 2004; Bullock et al., 2016; Dale et al., 2005; Fandos et al., 2023; Robledo-Arnuncio and Garcéia, 2007). Our results are thus consistent with a large panel of dispersal studies, and fat-tailed dispersal kernels seem to be a common property applicable to both our metapopulations (Klein et al., 2006; Petrovskii and Morozov, 2009). The comparison of our dispersal kernels with the one inferred by Lehnen et al. (2021) across a wide European metapopulation shows that our method is able to detect fine differences in kernel shape and leptokurtosis between two different metapopulations (see Fig. 4 and Fig. S5). The only parameter that significantly differed between the two distributions was the shape parameter of the Weibull distribution. The gametic kernel in Picardy shows a nearly gaussian mode around 20km, while in Thuringia the gametic kernel is more clearly partitioned between very short-scale dispersal (*<* 10km) and extreme long-distance dispersal events.

It remains difficult to evaluate if these differences could be due to power issues in population assignment or true variations in behaviour. Part of these differences could be caused by the lower power to infer the natal colony in Picardy, yet each metapopulation showed clearly different patterns even after integrating assignment uncertainty. Thuringia exhibits long dispersal distances that were not observed in Picardy (Fig. 2 and Fig. 4), a pattern that could be associated with the range expansion that recently occurred in this part of Germany (Jan et al., 2019). Finally, Picardy and Thuringia have in common the high proportion of intra-colonial paternities, which seems to be the main driver of the fine-scale genetic structure observed in both metapopulations.

Although long-distance dispersal events seem relatively rare, they remain important for gene flow and shape the leptokurtic dispersal kernel. Most strikingly, some of the actively dispersing males are involved in extreme long-distance mating events (see the map of mating dispersal movements in Fig. S1). Long-distance dispersal is often dependent on individual variation, with few individuals expressing extreme behaviours (Hawkes, 2009; Kesler et al., 2010), as was already observed for this species (Heymer, 1964). Interestingly, we observed that mating dispersal introduces variance in individual behaviours that define the leptokurtic dispersal kernel (Petrovskii and Morozov, 2009). Indeed we reported low median dispersal distances and the mean effective dispersal distances were strongly impacted by extreme values. Thus, individual variation in mating behaviours should be considered as a strong component of the dispersal kernel. Niggemann et al. (2012) reported that uncertainty in the dispersal kernel increases for long distances. Parentage assignment permits to assess this individual variation, but underestimation is expected. Some long-distance dispersal may remain cryptic due to the low number of fathers inferred (n = 34) and the finite size of the study area. Yet, our study species usually flies less than 40 km (Lehnen et al., 2021) and the sampling spatial scale was twice this distance.

### When sperm go further than individuals: mating dispersal decorrelates the flows of genes and individuals

At least 50% of males were moving temporarily outside their residence colony exclusively for mating, sometimes at very long distances, which suggests that mating dispersal is indeed a major component of gene flow in the lesser horseshoe bat, as already observed in a handful of philopatric mammal species. Longer mating than natal dispersal distances have been documented in kangaroo rats (*Dipodomys spectabilis*) and Spix’s disc-winged bat (*Thyroptera tricolor*), two species characterized by strong philopatry of both sexes (Buchalski et al., 2014; Winters and Waser, 2003).

Mating dispersal plays the same role in metazoans as gamete dispersal in aquatic (mainly marine) organisms and pollen dispersal in terrestrial plants: it allows gametes to move further than the individuals producing gametes, ultimately decoupling metapopulation dynamics and gene flow. This is an interesting context for enhancing our understanding of dispersal evolution. Available models and experiments highlight the role played by kin competition in favouring long-distance dispersal and thus fat-tailed dispersal distance distributions (Bitume et al., 2013; Ronce, 2007; Rousset and Gandon, 2002). The relatedness between mating partners decreases when mating dispersal movements increase in distance, so that their offspring are less likely to live in a kin environment. By contrast, mating dispersal has no influence on local density or the probability to reach empty habitat patches, two factors that are also thought to drive the evolution of dispersal distances (Bitume et al., 2013; Ronce, 2007). If kin competition is a potential key driver of mating dispersal, it may interact with other selective forces such as dispersal costs and inbreeding load. An attempt by Ravigné et al. (2006) to model the interactions between these different factors on the joint evolution of seed and pollen dispersal at different distances provides an example for the kind of information that could be gained from modelling dispersal in species that rely both on zygotic and gametic dispersal with different life-stages and behaviours involved. In this model, pollen dispersal distance depends on seed dispersal, with the outcome being conditional on the relative dispersal costs associated with short-versus long-distance and seed versus pollen dispersal. Finally, our indirect approach decomposing the total gametic dispersal into natal and mating dispersal components should be a useful addition to study complex dispersal patterns in animal species in which gene flow and movements of individuals are decorrelated, such as philopatric species.

### Methodological limitations for inferences of mating and gametic dispersal

Only a few paternities were considered as reliable after a stringent selection based on simulation results. This contrasts with the high number of mothers recovered for the same colonies (Zarzoso-Lacoste et al., 2018), indicating that the reduced number of inferred paternities is not linked to a lack of informativeness of the set of markers. An alternative explanation that must be considered is that many potential fathers were not sampled. In particular, only males haunting maternity colonies were sampled, i.e. males searching a priori the proximity of females. Some of the undetected males may avoid maternity colonies to form male colonies or to live solitarily. However this behaviour is known to diminish the mating success of fathers excluded from maternity colonies, therefore limiting their contribution to the mating dispersal kernel (Senior et al., 2005). More broadly in bat species, males seeking to access females generally reside in maternity colonies (Burland et al., 1999; Heckel and Von Helversen, 2003). Potential fathers are thus more likely to be among colonial fathers. Moreover, subsequent analyses will only be biased by this lack of potential fathers if unsampled fathers adopt a mating behaviour different from colonial fathers, which remains to be investigated. In contrast to other bat species (Burland et al., 1999; Petri et al., 1997; Rossiter et al., 2000), we found considerable reproductive skew amongst our colonial males, as only 17 males were responsible for all inferred paternities both in Picardy and Thuringia.

The two methods we employed for estimating mating dispersal distances (a categorical approach with COLONY and a Bayesian probabilistic approach with Masterbayes) yielded similar results, suggesting that COLONY estimates are robust. We also estimated natal dispersal distances by categorical genetic assignment to colonies, based on the selection of the best replicate among independent STRUCTURE runs. The reliability of this approach has already been assessed in a previous study, and genetic assignment with STRUCTURE remains robust to identify individual dispersers, compared to other methods (Dussex et al., 2016). After integrating the uncertainty in STRUCTURE analyses, we were able to detect a clear signal of natal philopatry. Finally, we combined mating and natal dispersal to estimate the gametic dispersal kernel based on a probabilistic modelling approach (Diefenbach et al., 2008; Williamson, 2002). Our results are in agreement with the dispersal kernel inferred by Lehnen et al. 2021.

### Male philopatry and its implications

Mating dispersal distance is generally higher in other, swarming bat species (Kerth and Morf, 2004; Rivers et al., 2005), compared to what we observed in the lesser horseshoe bat. This species has a sedentary lifestyle and low dispersal ability, and swarming has not yet been reported for this species (Schofield et al., 2022). This results in a strong genetic structure (Fig. S6) compared to other sedentary and forest specialist species such as the Bechstein’s bat (*Myotis bechsteinii*) (Kerth and Petit, 2005) or the brown long-eared bat (*Plecotus auritus*) (Burland et al., 1999). It is striking that genetic structure is stronger in the lesser horseshoe bat than the brown long-eared bat given that both are sedentary and that males and females share roosts in the latter one. However, in *Plecotus auritus*, males seek mating partners outside their colony (Burland and Worthington Wilmer, 2001).

Unexpectedly, we found 41 and 48% intra-colony paternities in two different metapopulations, which is a high percentage for a temperate bat (Kerth and Morf, 2004; Metheny et al., 2008) and also contrasts with swarming species (Burland et al., 1999; Kerth et al., 2003; Rivers et al., 2005). In the lesser horseshoe bat, pairs are formed between individuals of the same colony more often than by chance. If this trait is shared among rhinolophids, it could explain intra-lineage polygyny and more generally, mating among relatives (Rossiter et al., 2005). This also constrains dispersal distances and hence explains the fine-scale genetic structure we observe in this species. The scale at which isolation by distance can be detected is determined by the mean dispersal distance through its relationship with the second central moment of parent offspring distance, *σ* (Broquet and Petit, 2009; Rousset, 1997). Assuming isotropic dispersal, 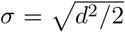, where *d* is the mean dispersal distance. In our case, this suggests that isolation by distance patterns could be detected beyond 12.4 and 15.7km in Picardy and Germany, respectively, which is consistent with the patterns we describe (Fig. S6). Interestingly, the estimates of dispersal distance based on general isolation by distance patterns (Lehnen et al., 2021) are close to the results we obtained through paternity analyses, supporting our assumption that there is no gene flow through females in this species. Our results should be relevant to a finer comprehension of why species with apparent sex-biased dispersal show nevertheless genetic structure at a fine scale.

## Conclusions

The population genetics approaches used here decomposed natal-dispersal and mating-dispersal components of the global gametic dispersal. This allowed to directly assess dispersal that would otherwise have remained cryptic if short-term movements for mating stayed undetected. Gene flow is intrinsically limited between lesser horseshoe colonies even when considering both natal and mating dispersal, despite long-distance dispersal events being more frequent than expected. We observed that variations in individuals’ mating behaviour shape a skewed dispersal kernel while promoting fine-scale genetic structure, with potential implications for genetic differentiation between populations. Hence, documenting these patterns is also important to help disentangle potential causes of dispersal evolution.

## Acknowledgments

We thank Gerald Kerth for sharing the Thuringia dataset ahead of publication. We thank every member of the INRAE DECOD lab (former ESE team) for their insightful discussions and kind support.

## Statement of authorship

E.J.P. conceived the study; S.J.P., D.Z.-L., L.L. and P.-L.J. produced the data, T.B. analysed the data; T.B. and E.J.P. wrote the manuscript. All authors commented on the findings and reviewed the manuscript.

## Data and code availability

The data reported in this article and the code to analyse the data are freely available in the INRAE data repository - DECOD collection (https://doi.org/10.57745/GBV4EA).

