## Supplementary Materials and Methods for "The geometry of gametic dispersal in a flying mammal, *Rhinolophus hipposideros*"

March 31, 2026

### Supplementary Information

#### Detection probability

COLONY needs information about the expected probability of a parent to be sampled. Detection probabilities for mothers in natural populations were around 0.9 according to Jan (2019). Because fathers were sampled in lower proportions in colonies, we estimated the theoretical detection probability based on their observed and expected proportions in sampled colonies. With a balanced theoretical sex-ratio (e.g. the expected probability of 0.5 to sample a male) and a male sampled proportion of only 0.28 in our dataset, the theoretical detection probability for males should be around 0.56 (observed/expected = 0.28/0.5). The probability that the parents were included among the candidates were thus set to 0.9 for mothers and to 0.56 for fathers.

#### Bayesian estimates of the mating dispersal kernel parameter with Masterbayes

Assuming a negative exponential distribution of mating distances (Hadfield et al., 2006; Nathan et al., 2012), the shape parameter $\beta$ of the dispersal kernel was estimated by a Monte Carlo Markov Chain (MCMC) procedure on a Bayesian model provided in the MasterBayes package (Hadfield et al., 2006). The mean dispersal distance of the exponential function of the dispersal kernel estimated with MasterBayes was the inverse of $\beta$. Paternity was a function of the distance between the father and the offspring. As for the COLONY analyses, the dataset was multigenerational. We excluded offspring as parents, and we excluded females as parents of a putative offspring when they were sampled in a different colony as the putative offspring. The convergence of the sampling chains was assessed based both on the trace and density plots and diagnostics implemented in the R package CODA (Plummer et al., 2006). Using MasterBayes (1,000,000 iterations, burnin of 20,000, thinning of 100), the shape parameter ($\beta$) for the negative exponential dispersal kernel has been estimated to 0.17 (IC95=[0.12; 0.22]) in France and 0.13 (IC95=[0.11; 0.15]) in Germany. Therefore, the estimated mean dispersal distance for mating was 6.0km (IC95=[4.47; 8.13]) in France and 7.73km (IC95=[6.66; 9.0]) in Germany. Markov chains were convergent.

#### COLONY Simulations

Contrary to the MasterBayes probabilistic approach, parentage assignment obtained with COLONY was categorical. The only paternities accepted were those of high certainty. Simulations were used both to find a statistic discriminant enough to exclude wrong paternities and to perform a power analysis of the categorical assignment method. Simulated genotype datasets were constructed from allele frequencies observed in each sampled population with the COLONY simulation module for Windows version 2.0.6.4 (Wang, 2013). Since kinships in the simulated population were all known, these data were useful to compare inferred paternities with expected ones. Error rates and allelic frequencies were those previously estimated from data. The simulation built parent-offspring relationships across a single generation, considering a population with a balanced sex-ratio (1:1). A theoretical random population of 4,000 parents and 3,892 offspring was simulated. The mating structure of the theoretical population considered that females were monogamous with only one offspring and males were polygamous with 0 to 3 offspring. Only 56% of simulated males and 90% of simulated females were sampled in order to follow the detection probabilities in natural populations. Each simulated dataset was replicated 50 times with different random seeds then analysed together to assess for a convergence on true fathers over multiple replicated runs. Different statistics were compared between two groups of assignments, whether a true or a wrong father was found for a given offspring. Investigated statistics were mean and variance of the probability value given by COLONY, and the assignment frequency of the best father (i.e. the proportion of runs where the best father was found). Besides, some incomplete genotypes (6/8 complete loci) were kept in the empirical dataset to retain more individuals in the analyses. Sensitivity analyses were conducted on the same theoretical population, in which the number of incomplete loci was parameterized as a genotype missing frequency of 2/16 missing alleles for 6/8 complete loci for 20% of the individuals, as in the empirical dataset. The distributions of assignments successes and errors were compared to detect variability in power due to partial genotypes. Successes were assignments of the true father when it was sampled and no assignment otherwise. It was a type I error when the wrong father was found and a type II error when a sampled father was not found.

#### Supplementary figures

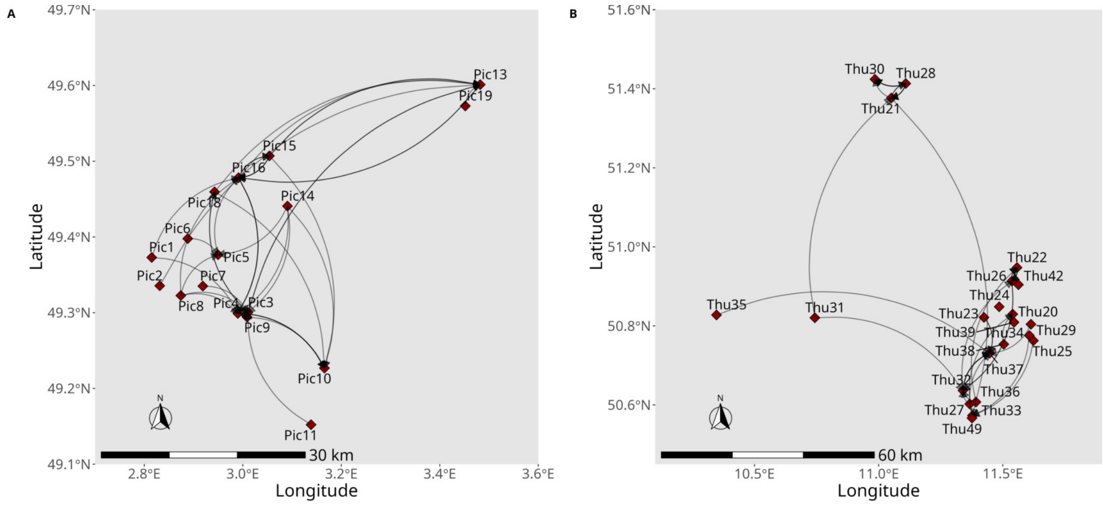


Figure S1: Geographical maps of sampled locations in France (A) and Germany (B). Grey arrows represent male mating dispersal events.

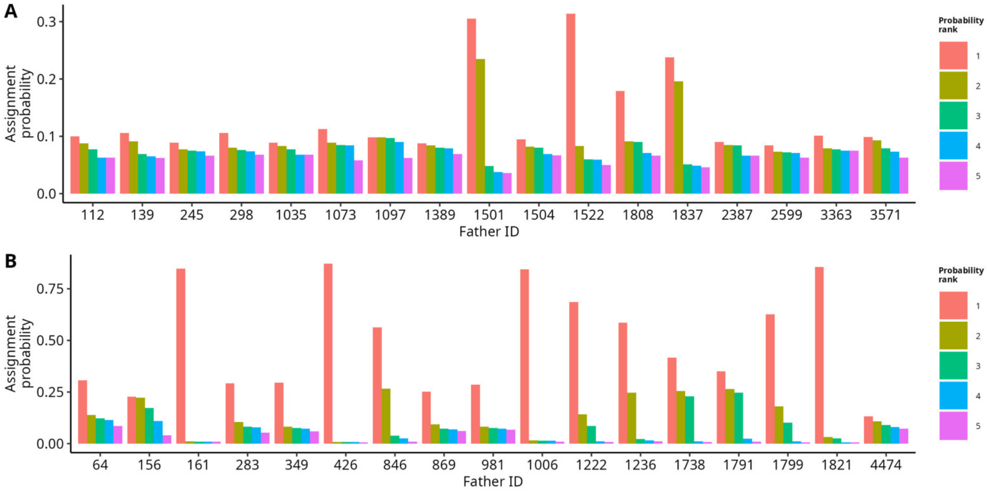


Figure S2: The five best natal colony assignment probabilities for all males inferred with COLONY in Picardy (A) and Thuringia (B) (n = 17 and 17, respectively). Assignment probabilities ordered by rank (i.e. ordered by decreasing probability value) for all fathers inferred in Picardy (A) and Thuringia (B). Only the five best probabilities are shown.

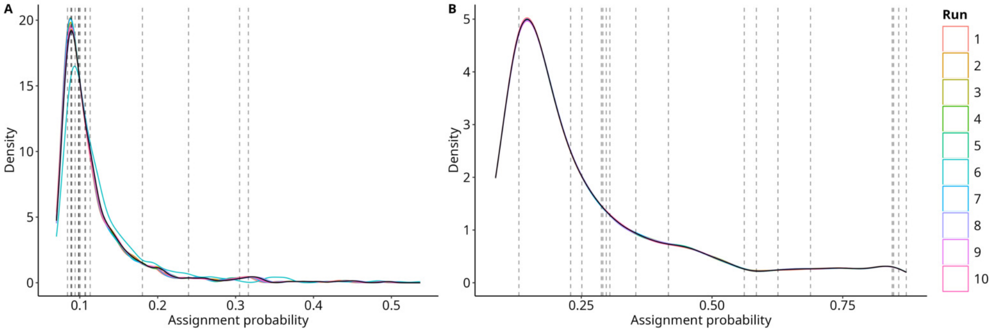


Figure S3: Density distributions of assignment probabilities for all adult males across ten independent MCMC sampling chains of STRUCTURE in Picardy (A) and Thuringia (B) (n = 578 and 683, respectively). Vertical dashed lines represent the best probability (rank 1) for fathers previously inferred with COLONY (n = 17 and 17, respectively).

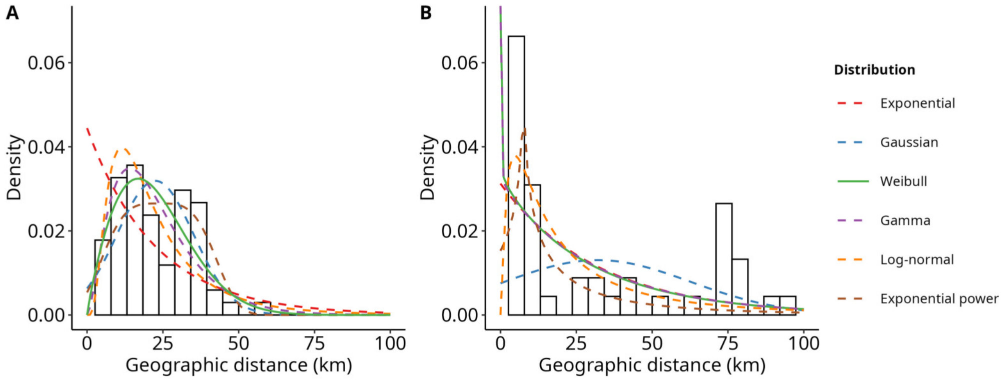


Figure S4: Competing models of gametic dispersal distance kernels fitted on strictly positive dispersal distances in Picardy (A) and Thuringia (B). Histograms show densities of empirical gametic dispersal distances (i.e. natal + mating dispersal). In each case, the selected model is indicated by the solid line and rejected models by dashed lines.

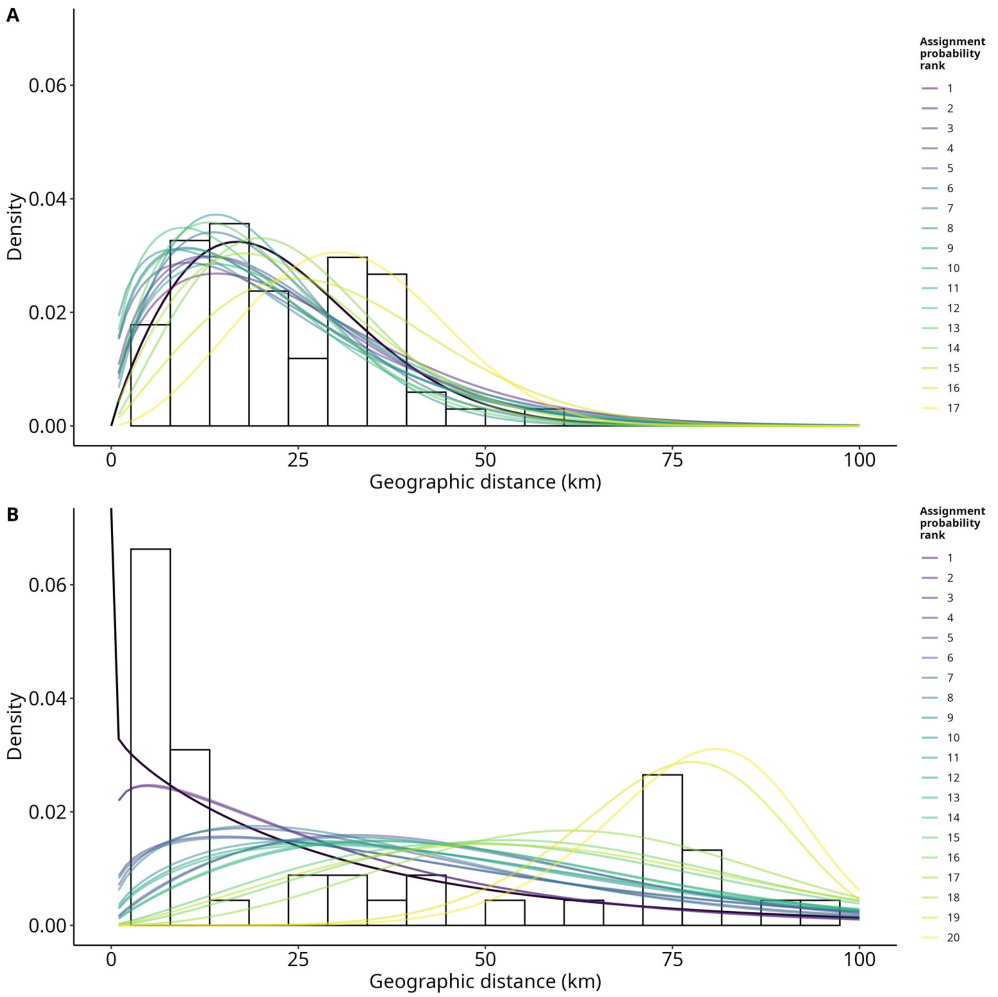


Figure S5: Integrating uncertainty in natal colony assignment in the gametic dispersal kernel in Picardy (A) and Thuringia (B). Comparison of gametic dispersal kernels estimated from positive dispersal distances between the natal colony of the father and the birth colony of the offspring, ordered by rank of the assignment probability, from the highest assignment probability to the lowest assignment probability inferred with STRUCTURE.

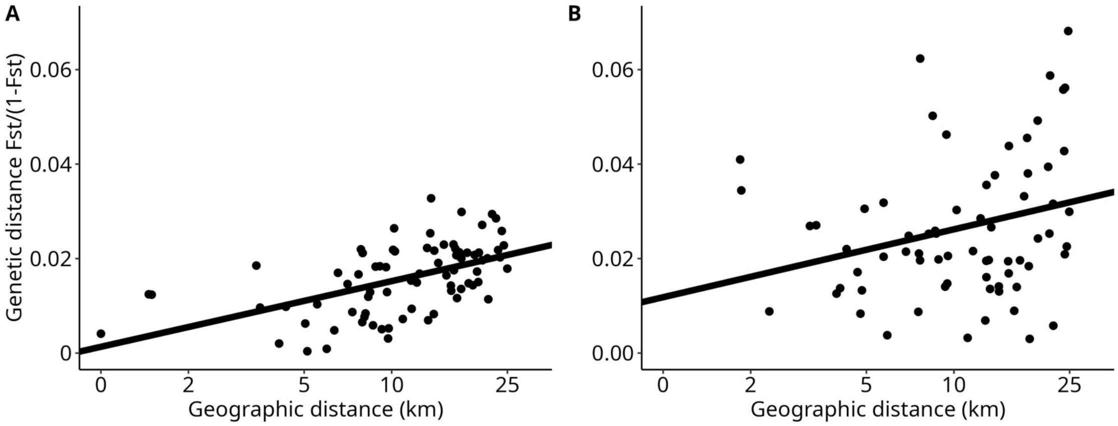


Figure S6: Fine-scale isolation-by-distance (IBD) pattern. We computed the relationship between the Euclidean distance in km (logarithmic scale) and the genetic distance FST/(1-FST) in females at a scale at which the pattern of IBD should be detected, based on theoretical arguments (see main text; Rousset, 1997). Pairwise FST estimated following Weir and Cockerham (1984). (A) Picardy ($y = 0.002 + 0.006x$; Mantel test, 1,000 permutations, $p < 0.001$). (B) Thuringia ($y = 0.01 + 0.006x$; Mantel test, 1,000 permutations, $p < 0.001$).

### Supplementary tables

Table S1: Genetic diversity for eight microsatellite markers in French and German populations (Picardy and Thuringia, respectively). Estimates of heterozygosity (Ho), gene diversity (Hs), and allelic richness were estimated with the R package hierfstat (Goudet and Jombart, 2022) and averaged across loci for each sampling site.

Table S2: Mean assignment frequency and standard deviation (SD) for each category of result in power simulations (simulated datasets). True fathers were known. (a) and (e) were correct results. (b) and (d) were type I errors, (c) was type II error. Expected frequencies were calculated under the hypothesis of 100% of correct results. Frequencies were calculated for 3,892 offspring and 30 runs.

| | | | | Picardy 8/8 loci | | Picardy 6/8 loci | | Thuringia 8/8 loci | | Thuringia 6/8 loci | |
|---|---|---|---|---|---|---|---|---|---|---|---|
| Results | | | Expected | Mean | SD | Mean | SD | Mean | SD | Mean | SD |
| (a) | Father sampled | True assignment | 0.56 | 0.07 | 0.01 | 0.03 | 0.004 | 0.025 | 0.004 | 0.011 | 0.002 |
| (b) | | False assignment | 0 | 0.04 | 0.01 | 0.04 | 0.01 | 0.034 | 0.008 | 0.036 | 0.008 |
| (c) | | No assignment | 0 | 0.45 | 0.02 | 0.49 | 0.01 | 0.51 | 0.011 | 0.5 | 0.01 |
| (d) | Father not sampled | False assignment | 0 | 0.05 | 0.01 | 0.04 | 0.01 | 0.024 | 0.006 | 0.028 | 0.007 |
| (e) | | No assignment | 0.44 | 0.39 | 0.01 | 0.4 | 0.01 | 0.403 | 0.006 | 0.421 | 0.007 |

Table S3: Paternities and maternities inferred in the French populations with COLONY.

Table S4: Paternities and maternities inferred in the German populations with COLONY.
