## Supplementary figures and images for "The geometry of gametic dispersal in a flying mammal, *Rhinolophus hipposideros*"

### Figure S1

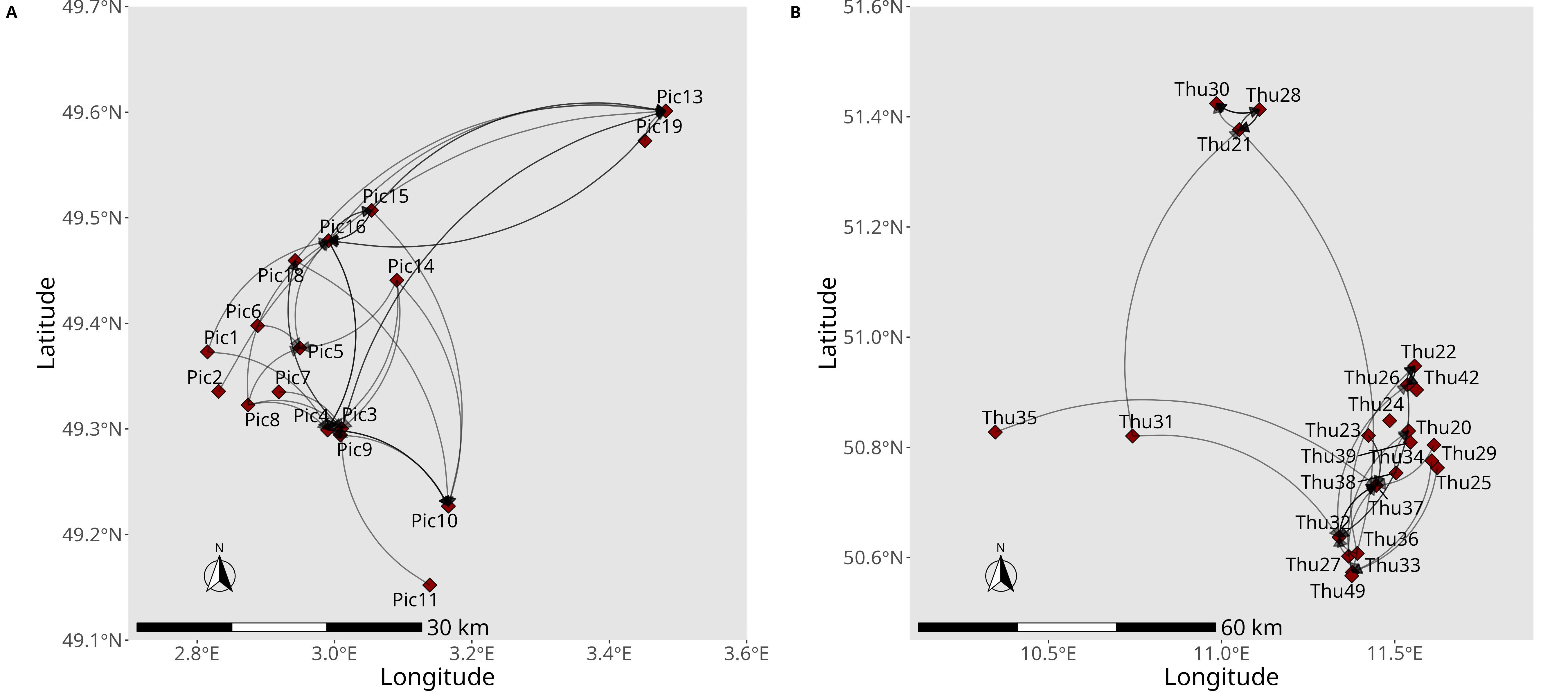

### Figure S2

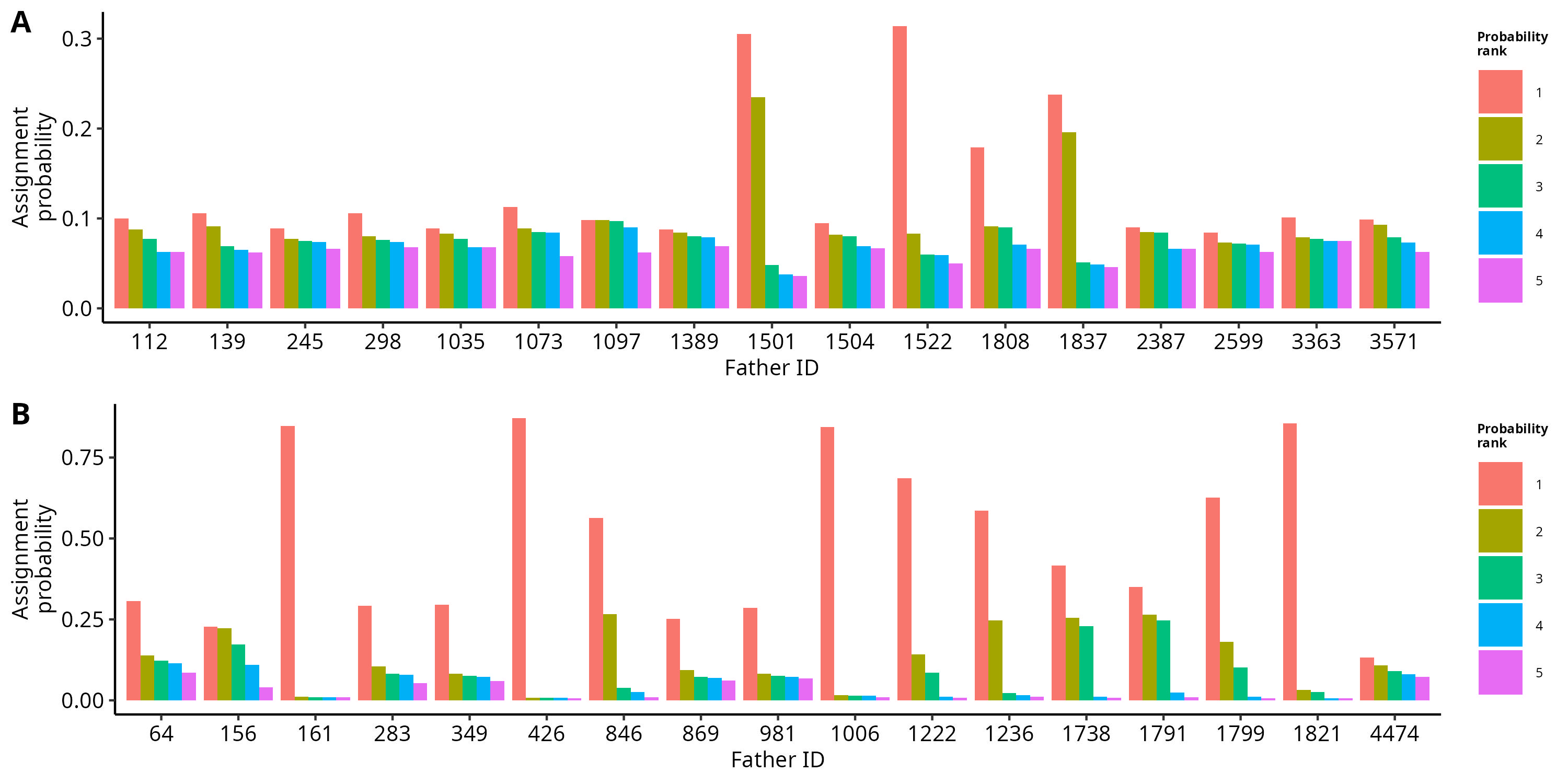

### Figure S3

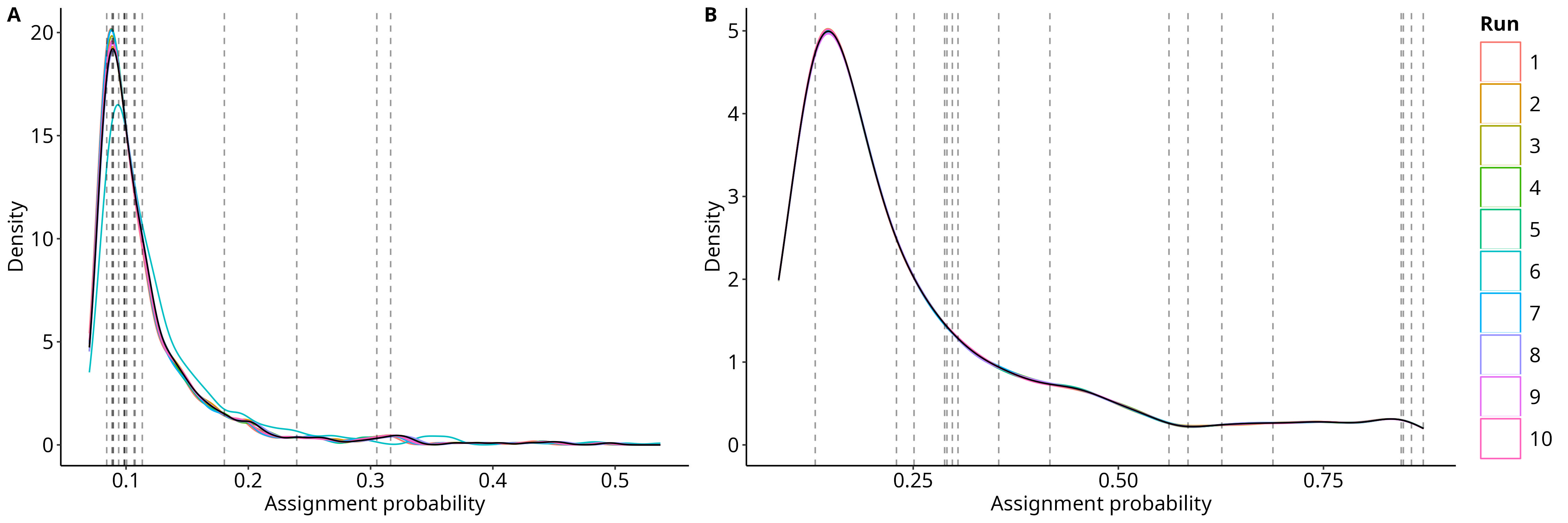

### Figure S4

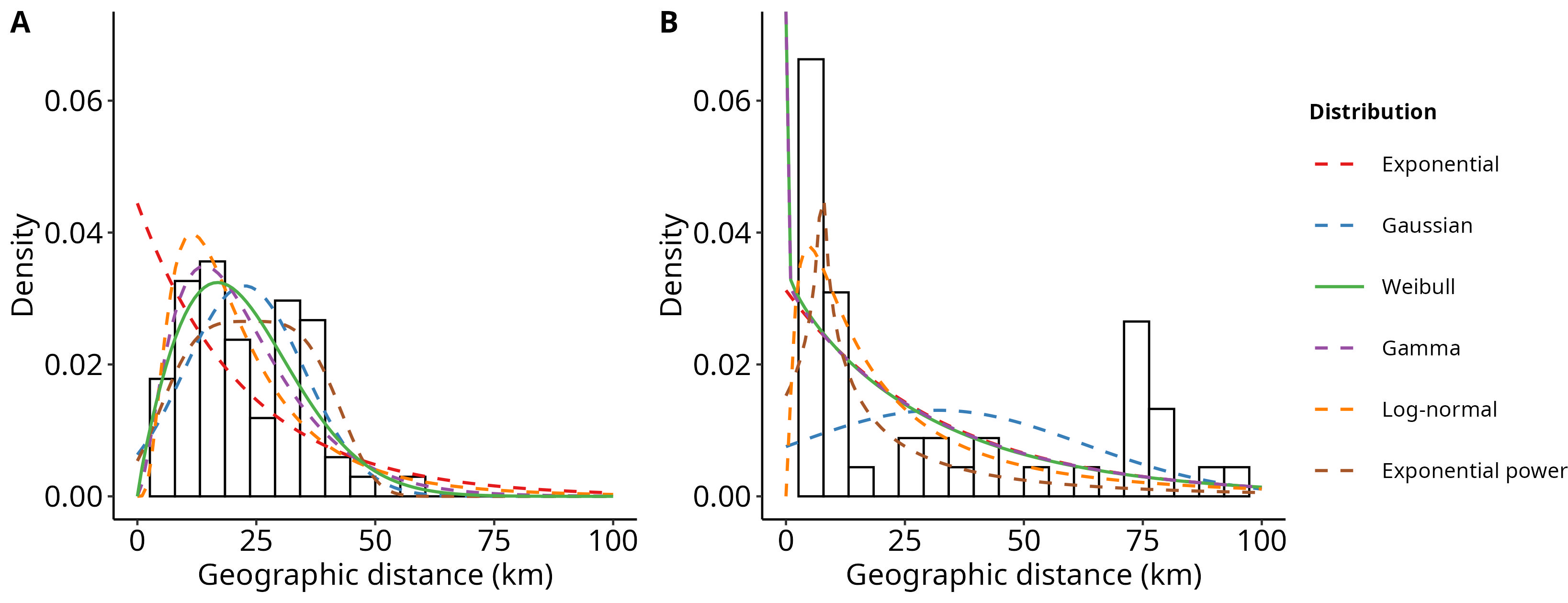

### Figure S5

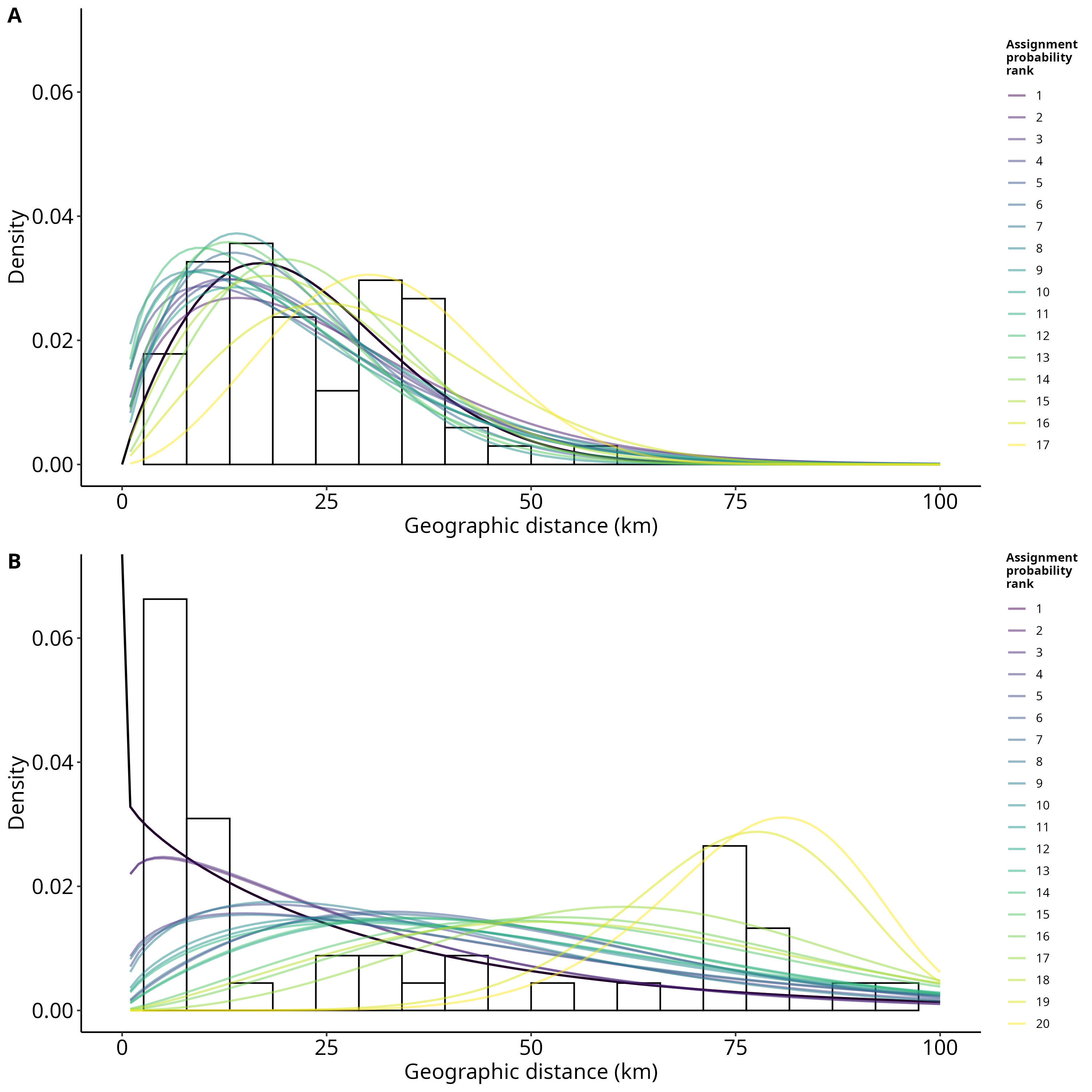

### Figure S6

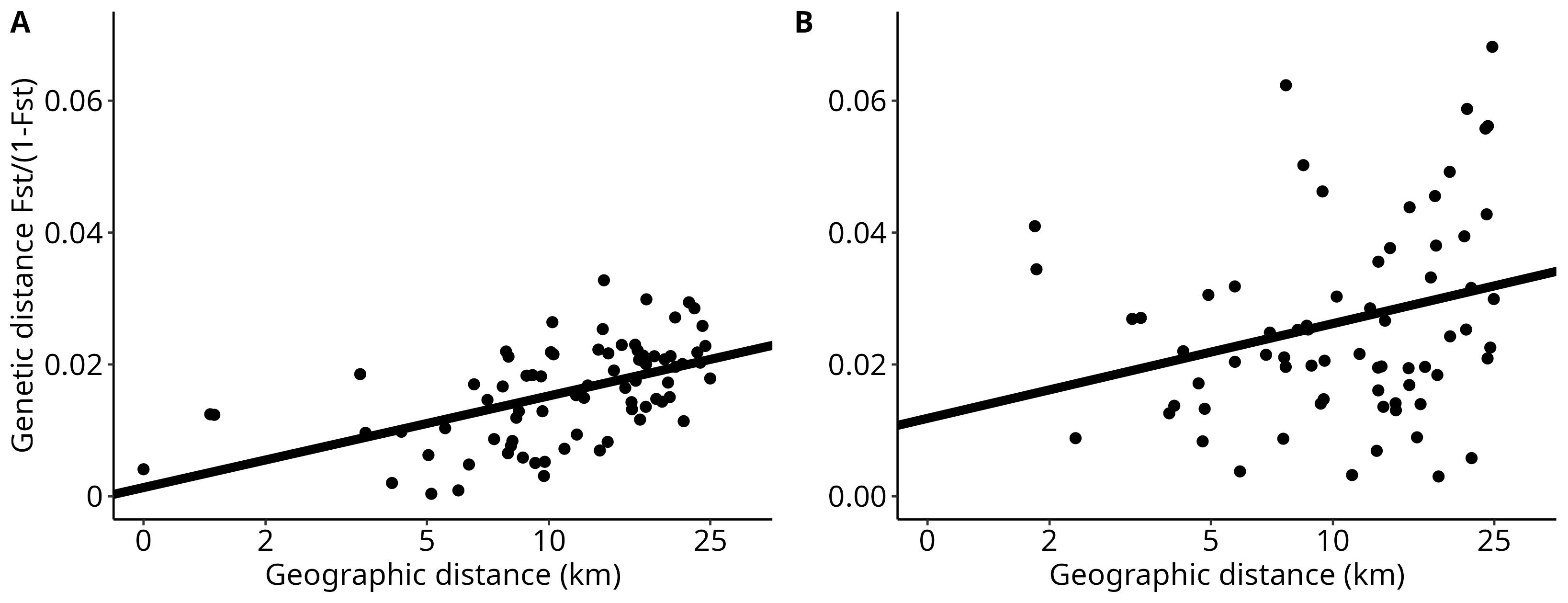
